# Genome-wide analysis of adolescent psychotic experiences shows genetic overlap with psychiatric disorders

**DOI:** 10.1101/265512

**Authors:** Oliver Pain, Frank Dudbridge, Alastair G. Cardno, Daniel Freeman, Yi Lu, Sebastian Lundstrom, Paul Lichtenstein, Angelica Ronald

## Abstract

This study aimed to test for overlap in genetic influences between psychotic experience traits shown by adolescents in the community, and clinically-recognized psychiatric disorders in adulthood, specifically schizophrenia, bipolar disorder, and major depression. The full spectra of psychotic experience domains, both in terms of their severity and type (positive, cognitive, and negative), were assessed using self- and parent-ratings in three European community samples aged 15-19 years (*Final N incl. siblings* = 6,297-10,098). A mega-genome-wide association study (mega-GWAS) for each psychotic experience domain was performed. SNP-heritability of each psychotic experience domain was estimated using genomic-relatedness-based restricted maximum-likelihood (GREML) and linkage disequilibrium-(LD-) score regression. Genetic overlap between specific psychotic experience domains and schizophrenia, bipolar disorder, and major depression was assessed using polygenic risk scoring (PRS) and LD-score regression. GREML returned SNP-heritability estimates of 3-9% for psychotic experience trait domains, with higher estimates for less skewed traits (Anhedonia, Cognitive Disorganization) than for more skewed traits (Paranoia and Hallucinations, Parent-rated Negative Symptoms). Mega-GWAS analysis identified one genome-wide significant association for Anhedonia within *IDO2* but which did not replicate in an independent sample. PRS analysis revealed that the schizophrenia PRS significantly predicted all adolescent psychotic experience trait domains (Paranoia and Hallucinations only in non-zero scorers). The major depression PRS significantly predicted Anhedonia and Parent-rated Negative Symptoms in adolescence. Psychotic experiences during adolescence in the community show additive genetic effects and partly share genetic influences with clinically-recognized psychiatric disorders, specifically schizophrenia and major depression.

## Text

### Introduction

Psychotic experiences, also referred to as psychotic-like experiences, are traits in the community that at the extreme resemble symptoms of psychotic disorders, such as schizophrenia. Based on principal component analyses, psychotic experiences can be separated into replicable and specific domains, such as positive, cognitive and negative domains [Ronald et al., 2014; Wigman et al., 2012]. Psychotic experiences are common in the general population, particularly during adolescence [Freeman, 2006; Ronald et al., 2014]. Evidence suggests that psychotic experiences are dimensional: they show varying degrees of severity and taxometric analyses support their continuous nature in adolescence [Ahmed et al., 2012; Daneluzzo et al., 2009; Taylor et al., 2016]. Adolescence is just prior to a peak time of onset for several psychiatric disorders, particularly schizophrenia, bipolar disorder, and major depression [Laursen et al., 2007]. Psychotic experiences predict many types of psychiatric disorders and suicidal ideation with significant odds ratios of 1.3-5.6 [Cederlöf et al., 2016; Fisher et al., 2013; Kelleher et al., 2012, 2014; McGrath et al., 2016; Werbeloff et al., 2012; Zammit et al., 2013].

Twin studies report modest heritability of psychotic experience domains with typically between a third and a half of the variance explained by genetic influences [Ericson et al., 2011; Hay et al., 2001; Hur et al., 2012; Linney et al., 2003; Polanczyk et al., 2010; Wigman et al., 2011; Zavos et al., 2014]. One study to date has reported SNP-heritability estimates ranging from 0-32%. These results suggest that common genetic variation plays a role in at least some types of adolescent psychotic experiences [Sieradzka et al., 2015], but larger studies are needed to offer accurate estimates.

The one previous genome-wide association study (GWAS) of adolescent psychotic experiences assigned 3,483 individuals to high- or low-scoring groups based on a measure of positive psychotic experiences that combined paranoia, hallucinations and delusions [Zammit et al., 2014]. No locus achieved genome-wide significance.

The Psychiatric Genomics Consortium 2 schizophrenia GWAS has been previously used twice to test for an association between genetic risk of schizophrenia and adolescent psychotic experiences (these followed one study using Psychiatric Genomics Consortium 1 schizophrenia GWAS results [Zammit et al., 2014]). The first of these studies used quantitative measures of individual psychotic experiences in adolescence in a sample of *N*=2,133-2,140 and reported no significant positive association between any of the psychotic experience domains and schizophrenia genetic risk [Sieradzka et al., 2014]. This study also reported no significant positive association with bipolar disorder genetic risk. A second study with *N*=3,676-5,444 tested for an association between schizophrenia genetic risk and dichotomous outcomes on several scales: a combined positive psychotic experiences scale (which included paranoia, hallucinations, delusions, and thought interference), a negative symptoms scale, a depressive disorder scale, and an anxiety disorder scale [Jones et al., 2016]. This second study reported a significant positive association between schizophrenia genetic risk and both high negative symptoms and high anxiety disorder. There was no association with the positive psychotic experiences or depressive disorder scales.

Our study aimed to test for overlap in genetic influences between specific psychotic experience domains in adolescence and psychiatric disorders-specifically, schizophrenia, bipolar disorder, and major depression. Four psychotic experience subscales were used, derived from principal component analysis: Paranoia and Hallucinations, Cognitive Disorganization, Anhedonia, and Parent-rated Negative Symptoms. We also present the largest GWAS to date of adolescent psychotic experiences using a community sample of 6,297-10,098 individuals (including siblings) from three European studies. As such our study also aimed to identify novel common genetic variants associated with specific psychotic experience domains and to estimate their SNP-heritability.

### Materials and Methods

#### Samples

Three European general population samples of comparable ages were included: TEDS (Twins Early Development Study) [Haworth et al., 2013] a community sample born in England and Wales (mean age 16.32 years); ALSPAC (Avon Longitudinal Study of Parents and Children) [Boyd et al., 2012] a cohort from the United Kingdom (mean age 16.76 years); CATSS (Child and Adolescent Twin Study in Sweden [Anckarsäter et al., 2011]) a twin cohort in Sweden (mean age 18.31 years). Sample descriptions and ethical consent information are available in Supplementary Note 1. Standard exclusion criteria for GWASs of behavioral and cognitive traits were used (Supplementary Table 1)[Docherty et al., 2010].

#### Measures

We established comparable quantitative scales of psychotic experience domains across samples. The subscales in the Specific Psychotic Experiences Questionnaire, used in TEDS, were the starting point [Ronald et al., 2014]. The other two samples had used similar self-report items that were mapped onto domains of paranoia, hallucinations, cognitive disorganization and anhedonia, and parent-rated negative symptoms. The original measures available within each sample are listed in Supplementary Note 2. The content of each psychotic experience domain used within each sample is given in Supplementary Tables 2-5. Cronbach’s alphas (measure of reliability) of each scale within each sample are listed in Table 1.

**Table I.**
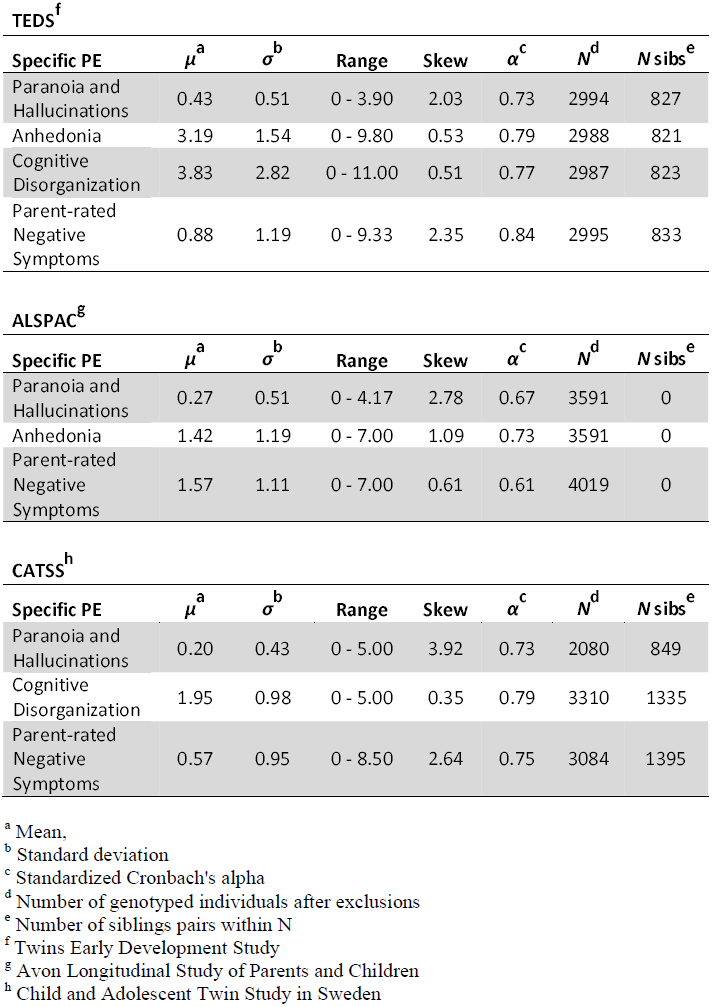
Descriptive statistics for psychotic experience domains in each sample. These figures are based on the individuals remaining after quality control and were used in all subsequent genetic analyses.

#### Phenotypic harmonization

Items were inspected to allow for harmonization of psychotic experience domains across samples via a two-stage process. First, two expert clinicians (AC and DF) matched items across samples that were capturing the same underlying construct (to ensure construct and content validity) based on their clinical knowledge and experience with psychotic experience measures. Second, the matched items within each sample were analyzed via principal components analysis, a psychometric approach used to determine whether items assessing each psychotic experience domain fell into individual components.

Individuals with >50% missingness for any psychotic experience domain were excluded from all analyses. The remaining missing phenotypic data was imputed using multiple imputation in R [Buuren and Groothuis-Oudshoorn, 2011]. Individual scores of the resulting psychotic experience subscales were calculated using sum scores. To ensure the equal contribution of each item to the sum score, item response values were rescaled to values between 0 and 1. The phenotypic correlations between specific psychotic experiences within each sample and in all samples combined are in Supplementary Table 6. Sum scores for each psychotic experience subscale were normalized using inverse rank-based normalization (data ties ranked randomly) and then standardized. The median correlation between a given scale before and after normalization was 0.92 (Supplementary Table 7), dependent on skew of the original scale. The normalized scores were then regressed against the following covariates: sex, age, age^2^, sex*age, sex*age^2^, study, and the top 8 principal components of ancestry. Principal components of ancestry were jointly calculated across all three samples using PLINK1.9 (https://www.cog-genomics.org/plink2) [Chang et al., 2015] based only on observed genetic variation.

#### Genotype imputation and quality control procedure

Genotypic data from each sample was imputed using the 1KG Phase 3 v5 to maximize genome-wide overlap between samples. After imputation, stringent variant and individual level quality control thresholds were applied in all samples before converting to hard-call genotypes (certainty threshold = 0.9) and merging the three datasets for combined analysis. For further information see Supplementary Note 3. Information on DNA collection and genotyping is given in Supplementary Note 4.

#### Estimation of SNP-heritability

SNP-heritability estimates were calculated within each sample and then meta-analyzed to provide meta-SNP-heritability estimates. The meta-analysis of SNP-heritability estimates approach was taken to avoid a downward bias due to possible genetic heterogeneity between samples. Inverse variance weighted meta-analysis was used. Secondary analysis of mega-SNP-heritability, estimated across samples simultaneously, was also performed to estimate the proportion of phenotypic variance that could be explained by the mean genetic effects across sample.

Two methods were used to estimate SNP-heritability: genomic-relatedness-matrix restricted maximum likelihood (GREML) in Genome-wide Complex Trait Analysis (GCTA), and linkage disequilibrium (LD)-score regression. Related individuals were included in both meta- and mega-GREML analyses [Zaitlen et al., 2013]. LD-score regression [Bulik-Sullivan et al., 2015] was also performed within (meta-) and across (mega-) samples. The effective sample size was used in LD-score regression analyses, thus matching the sample in the GWAS. Effective sample size was calculated as follows: (2*sample size)/(1+correlation between siblings) [Minica et al., 2014]. There was no evidence of confounding from genome-wide association results so the intercept was constrained to 1.

#### Mega-genome-wide association study

The three samples were combined to enable genome-wide association mega-analysis. Genome-wide association analysis of all four psychotic experience domains using related (i.e.monozygotic and dizygotic twin pairs) and unrelated individuals was performed in PLINK (http://pngu.mgh.harvard.edu/purcell/plink/) [Purcell et al., 2007]. Additional covariance arises from related individuals; this was accounted for using the method of generalizable estimating equations (GEE) [Minica et al., 2014; Minică et al., 2015] in R specifying an exchangeable correlation matrix.

#### Replication analysis of rs149957215 association with Anhedonia

After the discovery sample had been prepared and analysed for this project, an independent subsample of TEDS participants were genotyped. Of the newly genotyped individuals, 2,359 (incl. 635 MZ pairs) had reported on self-rated Anhedonia. This independent TEDS sample was used as a replication sample for the validation of the genome-wide significant association between rs149957215 and Anhedonia identified by the mega-GWAS in this study. rs149957215 genotypes were imputed (MACH r^2^ = 0.93) using the haplotype reference consortium data via the Sanger imputation server [McCarthy et al., 2016]. The genotypic data was converted to hard-call format (certainty threshold of 0.9) and analyzed in PLINK using the same GEE method to account for relatives. This replication analysis had a power of 0.86 to detect an association of the same magnitude (*r*^2^ = 0.47%) at nominal significance.

#### Gene-based association analysis

Two gene-based association analyses were performed. The first aggregates SNP associations within specific gene regions using the MAGMA program [de Leeuw et al., 2015]. The second analysis used PrediXcan [Gamazon et al., 2015] to predict frontal cortex gene expression differences using genotypic data. Further details can be found in Supplementary Note 5.

#### Genetic association between psychotic experience domains and psychiatric disorders

Using the software PRSice [Euesden et al., 2015], polygenic risk scores (PRSs) of schizophrenia, bipolar disorder, and major depression were calculated in the adolescent sample using the log of the odds ratios from the latest Psychiatric Genomics Consortium GWAS of schizophrenia (PGC2) [Ripke et al., 2014], bipolar disorder [PGC Bipolar Disorder Working Group, 2011] and major depression [Ripke et al., 2013]. LD was controlled for using LD-based clumping with an *r*^2^-cutoff of 0.1 within a 250-kb window. For each individual, scores were generated using SNPs with the following *p*-value thresholds (*p*T): 0.001, 0.01, 0.05, 0.1, 0.2, 0.3, 0.4, and 0.5. Linear regressions were performed in R and GEE was used to account for related individuals. Quantile plots were created to further examine the relationships. If there was evidence of a non-linear relationship from the quantile plots, the non-linear relationship was formalized by performing linear regression in a subset of individuals.

To estimate the genetic covariance between psychotic experience domains and schizophrenia, bipolar disorder, and major depression, both LD-score regression and AVENGEME (Additive Variance Explained and Number of Genetic Effects Method of Estimation) were used [Bulik-Sullivan et al., 2015; Palla and Dudbridge, 2015]. AVENGEME uses the results of polygenic risk score analyses across multiple significance thresholds to estimate the model parameters including the genetic covariance. AVENGEME estimates 95% confidence intervals using profile likelihood method. There was no evidence of confounding or sample overlap in the mega-GWAS summary statistics, as such the heritability-intercept was constrained to 1 and the genetic covariance intercept was set to 0 in LD-score regression. To improve the accuracy of the estimates of genetic covariance derived from the AVENGEME analysis, the SNP-heritability of liability for schizophrenia, bipolar disorder, and major depression were constrained to the LD-score regression estimates of SNP-heritability (see Supplementary Table 8).

### Results

Table 1 shows the descriptive statistics of psychotic experience domains split by sample. After phenotypic harmonization and quality control, the final sample sizes (including siblings) for all subsequent analyses were 8,665 for Paranoia and Hallucinations, 6,579 for Anhedonia, 6,297 for Cognitive Disorganization, and 10,098 for Parent-rated Negative Symptoms.

SNP-heritability estimates from meta-GREML and meta-LD-score regression were between 2.8-8.8% and 6.6-21.5% respectively (Table 2). Results from secondary analysis of SNP-heritability using the mega-analysis approach are given in Supplementary Table 9.

**Table II.** SNP-heritability estimates for psychotic experience domains.

| Specific PE | meta-GREML-SC |  | meta-LDSC |  |
| --- | --- | --- | --- | --- |
| | SNP-$h^2$<sup>a</sup> | SE | SNP-$h^2$<sup>a</sup> | SE |
| Paranoia and Hallucinations | 0.028 | -0.050 | 0.066 | 0.068 |
| Anhedonia | 0.088 | 0.055 | 0.204 | 0.078 |
| Cognitive Disorganisation | 0.059 | 0.063 | 0.215 | 0.085 |
| Parent-rated Negative Symptoms | 0.059 | 0.045 | 0.119 | 0.061 |

<sup>a</sup> Phenotypic variance explained by tagged common genetic variation

The mega-GWAS identified no genome-wide significant variation for the Paranoia and Hallucinations, Cognitive Disorganization, or parent-rated Negative Symptoms domains. The mega-GWAS of Anhedonia identified one SNP (rs149957215) associated at genome-wide significance (*p* = 3.76x10^-8^) within a gene called indoleamine 2,3-dioxygenase 2 (*IDO2***) (**Supplementary Figure 1). This SNP was imputed in both samples included in the Anhedonia GWAS (TEDS and ALSPAC) with an average imputation quality (INFO) of 0.87 and minor allele frequency of 0.015. The minor allele frequency was similar in both TEDS (0.017) and ALSPAC (0.013). Due to limited LD with rs149957215, neighboring genetic variation showed no significant evidence of association. This association between rs149957215 and Anhedonia did not replicate in the independent TEDS replication sample, showing a non-significant association (*p*=0.81) in the opposite direction. Several loci achieved suggestive significance (*p*<1x10^-5^) across the four psychotic experience domains (Supplementary Table 10, Supplementary Figures 2-5). There was no evidence of confounding with lambdas of 0.99–1.01 and LD-score regression intercept of 1.00 in all analyses (Supplementary Figure 6).

Regional gene-based tests identified no gene that was significantly associated with any psychotic experience domain after Bonferroni correction for multiple testing. The top ten most associated genes for each psychotic experience domain are listed in Supplementary Table 11.

Analysis of predicted frontal cortex gene expression associated with psychotic experience domains showed *HACD2* as significantly differentially expressed for Cognitive Disorganization (Bonferroni corrected *p* = 6.83x10^-4^). The top ten genes showing differential expression for each psychotic experience domain are listed in Supplementary Table 12.

The schizophrenia PRS significantly and positively predicted Anhedonia (*p* = 0.030 at *p*T = 0.10), Cognitive Disorganization (*p* = 0.035 at *p*T = 0.01) and Parent-rated Negative Symptoms (*p* = 5.41x10^-3^ at *p*T = 0.05) (Table 3; Figure 1). The bipolar disorder PRS significantly and negatively predicted Paranoia and Hallucinations only (*p* = 2.47x10^-3^ at *p*T= 0.010) (Table 3; Figure 1). The major depression PRS significantly and positively predicted Anhedonia (*p* = 0.010 at *p*T 0.5) and Parent-rated Negative Symptoms (*p* = 8.29x10^-3^ at *p*T = 0.001) (Table 3; Figure 1). Supplementary Tables 13-15 and Supplementary Figures 7-9 show the full results of these analyses. Logistic regression comparing PRSs in low and high psychotic experience domain groups (defined as bottom and top 25% of raw psychotic experience sum scores) were congruent with linear analyses (Supplementary Table 16, Supplementary Figure 10).

**Figure 1.**
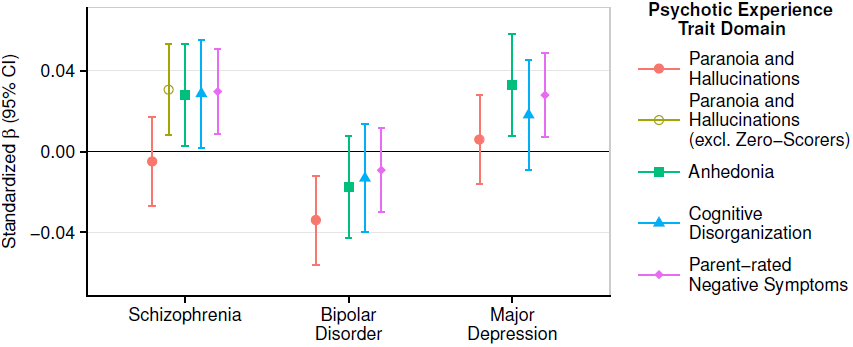
Polygenic risk scores for schizophrenia, bipolar disorder, and major depression predict adolescent psychotic experience domains. This figure shows results for polygenic risk scores at the most predictive *p*-value threshold for each trait. Error bars are 95% confidence intervals (95% CI).

**Table III.** Schizophrenia, bipolar disorder, and major depression polygenic risk scores predicting psychotic experience domains in adolescents. This table shows results for polygenic risk scores at the most predictive *p*-value threshold for each trait.

| **Specific PE** | **$\beta$** | **SE** | **$p$** | **$r^2$** | **$pT^a$** |
| --- | --- | --- | --- | --- | --- |
| Paranoia and Hallucinations | -0.005 | 0.011 | 0.664 | 0.002% | 0.200 |
| Paranoia and Hallucinations<br>excl. zero-scorers | 0.031 | 0.012 | **7.90E-03** | 0.094% | 0.001 |
| Anhedonia | 0.028 | 0.013 | **0.030** | 0.079% | 0.100 |
| Cognitive Disorganisation | 0.029 | 0.014 | **0.035** | 0.083% | 0.010 |
| Parent-rated Negative<br>Symptoms | 0.030 | 0.011 | **5.41E-03** | 0.088% | 0.050 |

***Bipolar Disorder***
| **Specific PE** | **$\beta$** | **SE** | **$p$** | **$r^2$** | **$pT^a$** |
| --- | --- | --- | --- | --- | --- |
| Paranoia and Hallucinations | -0.034 | 0.011 | **2.47E-03** | 0.115% | 0.010 |
| Anhedonia | -0.017 | 0.013 | 0.178 | 0.030% | 0.100 |
| Cognitive Disorganisation | -0.013 | 0.014 | 0.333 | 0.017% | 0.001 |
| Parent-rated Negative<br>Symptoms | -0.009 | 0.011 | 0.388 | 0.008% | 0.500 |

***Major Depression***
| **Specific PE** | **$\beta$** | **SE** | **$p$** | **$r^2$** | **$pT^a$** |
| --- | --- | --- | --- | --- | --- |
| Paranoia and Hallucinations | 0.006 | 0.011 | 0.589 | 0.004% | 0.100 |
| Anhedonia | 0.033 | 0.013 | **0.010** | 0.109% | 0.500 |
| Cognitive Disorganisation | 0.018 | 0.014 | 0.189 | 0.033% | 0.050 |
| Parent-rated Negative<br>Symptoms | 0.028 | 0.011 | **8.29E-03** | 0.078% | 0.001 |

<sup>a</sup> p-value threshold used to select when calculating polygenic risk scores

Quantile plots showing the mean PRS within subsets of the PE distributions highlighted one non-linear relationship between the schizophrenia PRS and Paranoia and Hallucinations, with the point of inflection at the median (Supplementary Figure 11). The majority of individuals (81%) below the median had a raw score of zero. A single post-hoc analysis was performed to formalize the presence of this non-linear relationship. Post-hoc removal of individuals with a raw Paranoia and Hallucinations score of zero led to the schizophrenia PRS positively predicting Paranoia and Hallucinations (*p* = 7.90x10^-3^ at *p*T = 0.001) (Table 3; Figure 1; Supplementary Table 13). Logistic regression comparing low and high groups of non-zero scoring individuals supported these findings (Supplementary Table 16, Supplementary Figure 10).

Table 4 presents the AVENGEME estimates of genetic covariance, which were highly congruent with the PRS analysis results. AVENGEME pools evidence across *p*-value thresholds tested from PRS analysis. As such, even when there is consistent but non-significant evidence of association at individual *p*-value thresholds, AVENGEME genetic covariance estimates can be significant. Consequently, there were two significant results that were not shown by the PRS analyses; between the Anhedonia psychotic experience domain and bipolar disorder (genetic covariance = -0.022, 95%CI = -0.041 - -0.002), and the Cognitive Disorganization psychotic experience domain and major depression (genetic covariance = 0.033, 95%CI = 0.005 – 0.062).

**Table IV.** Genetic covariance between each psychotic experience domain and schizophrenia, bipolar disorder, and major depression.

**Schizophrenia**
| Specific PE | AVENGEME |  |  | LDSC |  |  |
| --- | --- | --- | --- | --- | --- | --- |
|  | *cov(G)*<sup>a</sup> | Low<br>95% CI | High<br>95% CI | *cov(G)*<sup>a</sup> | SE | *p* |
| Paranoia and Hallucinations | -0.002 | -0.013 | 0.008 | -0.003 | 0.014 | 0.844 |
| Paranoia and Hallucinations<br>excl. zero-scorers | **0.019** | 0.011 | 0.029 | NA | NA | NA |
| Anhedonia | **0.024** | 0.013 | 0.038 | 0.021 | 0.016 | 0.184 |
| Cognitive Disorganisation | **0.025** | 0.013 | 0.040 | **0.054** | 0.018 | 3.52E-03 |
| Parent-rated Negative<br>Symptoms | **0.025** | 0.015 | 0.036 | **0.047** | 0.014 | 5.57E-04 |

**Bipolar Disorder**
| Specific PE | AVENGEME |  |  | LDSC |  |  |
| --- | --- | --- | --- | --- | --- | --- |
|  | *cov(G)*<sup>a</sup> | Low<br>95% CI | High<br>95% CI | *cov(G)*<sup>a</sup> | SE | *p* |
| Paranoia and Hallucinations | **-0.032** | -0.048 | -0.018 | **-0.045** | 0.022 | 0.039 |
| Anhedonia | **-0.022** | -0.041 | -0.002 | 0.008 | 0.028 | 0.767 |
| Cognitive Disorganisation | 0.008 | -0.013 | 0.029 | 0.021 | 0.028 | 0.451 |
| Parent-rated Negative<br>Symptoms | -0.010 | -0.026 | 0.006 | 0.009 | 0.021 | 0.669 |

**Major Depressive Disorder**
| Specific PE | AVENGEME |  |  | LDSC |  |  |
| --- | --- | --- | --- | --- | --- | --- |
|  | *cov(G)*<sup>a</sup> | Low<br>95% CI | High<br>95% CI | *cov(G)*<sup>a</sup> | SE | *p* |
| Paranoia and Hallucinations | 0.004 | -0.018 | 0.026 | 0.013 | 0.020 | 0.523 |
| Anhedonia | **0.065** | 0.037 | 0.094 | **0.061** | 0.023 | 7.50E-03 |
| Cognitive Disorganisation | **0.033** | 0.005 | 0.062 | 0.015 | 0.028 | 0.594 |
| Parent-rated Negative<br>Symptoms | **0.023** | 0.010 | 0.042 | 0.031 | 0.020 | 0.119 |

<sup>a</sup> Genetic covariance

Table 4 presents estimates of genetic covariance from LD-score regression. LD-score regression mirrored over half of the significant associations shown between the equivalent polygenic risk scores and psychotic experience domains in Table 3. The genetic covariance between the schizophrenia PRS and non-zero scorers on Paranoia and Hallucinations could not be estimated because the genome-wide association analysis included zero scorers. Unlike for the equivalent results in Table 3, the genetic covariance between schizophrenia and anhedonia psychotic experience domain, and between major depression and parent-rated negative symptoms domain, were not significant.

### Discussion

A genetic relationship was identified between clinical schizophrenia and positive, cognitive, and negative psychotic experience trait domains in adolescence. A higher genetic risk for schizophrenia significantly predicted adolescents having more cognitive disorganization, anhedonia, and parent-rated negative symptoms, as well as more paranoia and hallucinations (this last finding was in the non-zero scorers). Thus our findings suggest that psychotic experience trait domains in adolescence comprise partly of the genetically-influenced phenotypic manifestation of schizophrenia. Furthermore, higher genetic risk for major depression significantly predicted having more self-rated anhedonia and parent-rated negative symptoms as a teenager. Our results discredit the hypothesis that psychotic experience trait domains in the general population are epiphenomena that do not share biological pathways with clinically-recognized psychiatric disorders.

Genetic risk for schizophrenia and major depression in adulthood predicted 0.08– 0.09% variance in psychotic experience domains in adolescence. The effect sizes are likely downward biased because of the conservative approaches taken here to align different samples and to handle skewed variables. Nevertheless, comparable effect sizes have been reported in equivalent studies on other phenotypes, for example, linking social communication traits to genetic liability for autism [St Pourcain et al., 2017]. Large effect sizes were not expected. The upper limit of prediction is the degree to which polygenic risk scores for schizophrenia and major depression predict themselves (that is, their own phenotype) in an independent sample, which is ~18.4% for schizophrenia [Ripke et al., 2014] and ~0.6% for major depression [Ripke et al., 2013]. It is known from epidemiological studies that the magnitude of phenotypic association between psychotic experience domains and schizophrenia is modest [McGrath et al., 2016; Zammit et al., 2013] and far more people report psychotic experiences in adolescence than develop disorders such as schizophrenia. Furthermore, there is considerable heterogeneity in schizophrenia and depression in terms of age of onset and symptom presentation. The associations were likely to be modest given their cross-phenotype and cross-age nature. The genetic association between psychiatric disorders and psychotic experience traits in the community may increase with age or with longitudinal assessments.

A notable result was that paranoia and hallucinations during adolescence are associated with schizophrenia common genetic risk if individuals report at least some degree of paranoia or hallucinations, that is, the association was only present in the non-zero scorers. Individuals reporting no paranoia and hallucinations can exist anywhere on the schizophrenia genetic liability spectrum. Previous studies did not find a genetic association between schizophrenia and positive psychotic experiences [Jones et al., 2016; Zammit et al., 2014; Sieradzka et al., 2014]. The use of quantitative traits and a large sample allowed the identification of this non-linear effect here. An explanation for the non-linear effect may lie in the variable age of onset of paranoia and hallucinations. Our study was focused on psychotic experiences in mid to late adolescence. It is predicted that stronger positive associations between genetic risk for schizophrenia and positive psychotic experiences will be found in samples assessed over a longer time frame, when anyone who is going to have paranoia and hallucinations has manifested them.

The significant and positive genetic association between a self-rated anhedonia trait measure in adolescence and major depression in adulthood concurs with a previous report showing that subclinical depressive symptoms (including anhedonia) phenotypically predict major depressive episodes in adulthood [Pine et al., 1999]. Anhedonia is present as a symptom of both schizophrenia and depression in psychiatric diagnoses, and our research shows that as a trait dimension in adolescence it shares common genetic underpinnings with both schizophrenia and depression.

The significant negative association between paranoia and hallucinations and bipolar disorder genetic risk also deserves discussion. Previous research reported the presence of paranoia, hallucinations, and delusions prior to the onset of bipolar disorder [McGrath et al., 2016]. Our results suggest that the manifestation of paranoia and hallucinations as traits during mid-adolescence do not share variance with known common genetic risk associated with diagnosed bipolar disorder.

Although in many cases the SNP-heritability estimates were not significantly non-zero, the point estimates of SNP-heritability indicate that common genetic variation influences psychotic experience domains during adolescence. This concurs with results from twin studies reporting significant twin heritability estimates [Zavos et al., 2014].

The SNP that achieved genome-wide significance in the mega-GWAS for anhedonia lies within the protein-coding gene *IDO2*. IDO2 is a key enzyme in the regulation of the kynurenine pathway, which upon stimulation by proinflammatory cytokines, converts tryptophan into kynurenine. It has been reported that increased metabolism of tryptophan to kynurenine is associated with increased depressive symptoms via the increased production of cytotoxic kynurenine metabolites [Wichers et al., 2005; Dantzer et al., 2011; Myint et al., 2007]. In fact, a previous study has reported a significant correlation between kynurenine production and anhedonia in an adolescent sample [Gabbay et al., 2012]. These previous studies suggest the association between *IDO2* and anhedonia is plausible. However, this association should be interpreted with caution as rs149957215 was imputed in all three samples, has a low minor allele frequency of 0.013, and appears to be uncorrelated with surrounding common genetic variation. Furthermore, this association failed to replicate in a sufficiently powered independent sample.

Some methodological details deserve consideration. A key strength of the study is the derivation of psychometrically-sound quantitative individual psychotic experience domains: these were derived using principal component analysis and have content validity. It is noted that our terminology of psychotic experiences, like others [Stefanis et al., 2002; Wigman et al., 2011; Alemany et al., 2013], includes positive, cognitive and negative psychotic experiences, for the reason that at the extreme these experiences are characteristic of clinical psychotic disorders. Greater power was achieved compared to past research through combining independent samples. At the same time, slight variations in measure items across samples impacts the amount of phenotypic variance that can be explained by common genetic variation. For the more skewed psychotic experience domains with larger numbers of tied individuals, the process of randomly ranking tied individuals during normalization was essential but will have introduced noise and thus downward biased SNP-heritability estimates. This study has identified significant common genetic covariance between several psychotic experiences and psychiatric disorders. It remains possible that the genetic covariance estimates could be partly explained by the presence of adolescents with diagnosed relatives.If an adolescent has a relative with schizophrenia for example, the adolescent may have increased psychotic experiences due to their shared environment with the relative. However, twin and adoption studies of schizophrenia and psychotic experiences do not support this mode of transmission. Finally, each psychotic experience domain was viewed as a phenotype of interest in its own right, and therefore view this study as a series of self-contained GWAS and genetic covariance studies, rather than a single exploratory study across traits. Therefore we did not formally correct for multiple testing of traits, similar to other studies of this nature [Consortium, 2007; Cross-Disorder Group of the Psychiatric Genomics Consortium,2013]. Note however that nearly all of our associations would remain significant after correction for testing four PE domains, while the association of rs149957215 with Anhedonia (*p*=3.76e-8) did not replicate.

Collectively, these findings reveal novel evidence for some shared common genetic etiology between psychotic experience domains in mid to late adolescence and clinically-recognized psychiatric disorders in adulthood. Evidence is accruing that psychotic experiences manifested prior to adulthood form part of a wider phenotype or prodrome related to psychiatric disorders such as schizophrenia and depression. This study joins the existing evidence that individuals with a family history of schizophrenia score higher on psychotic experience scales [Zavos et al., 2014; Jeppesen et al., 2014], psychotic experiences are associated with the same environmental risk factors as schizophrenia [Linscott and Van Os, 2013], psychotic experiences are on a phenotypic and etiological continuum across the severity continuum [Zavos et al., 2014; Taylor et al., 2016], and psychotic experiences predict later psychiatric disorders [McGrath et al., 2016; Fisher et al., 2013; Zammit et al., 2013; Cederlöf et al., 2016; Kelleher et al., 2014; Werbeloff et al., 2012; Kelleher et al., 2012]. The next step is to consider how psychotic experiences can be harnessed in a practical sense as a (small effect size) red flag for risk in early intervention and prevention strategies. Similar to family history, psychotic experiences might be a useful heuristic even if the majority of individuals with psychotic experiences will remain unaffected by psychiatric disorders.

## Acknowledgments

We thank the participants of TEDS, ALSPAC and CATSS, and their research teams, which include interviewers, computer and laboratory technicians, clerical workers, research scientists, volunteers, managers, receptionists and nurses. We also thank the Psychiatric Genomics Consortium for the publically available GWAS summary statistics employed here. We thank Robert Plomin, George Davey Smith, and Camelia Minica for her script for performing GWAS with GEE to account for relatives.

The TEDS study of psychotic experiences was funded by Medical Research Council grant G1100559 to AR and a Wellcome Trust ISSF grant to AR. TEDS is funded by Medical Research Council grant G0901245, and G0500079 to Robert Plomin. OP was funded by a Bloomsbury PhD studentship. DF is supported by an NIHR Research Professorship. The UK Medical Research Council and Wellcome (Grant ref: 102215/2/13/2) and the University of Bristol provide core support for ALSPAC. ALSPAC GWAS data was generated by Sample Logistics and Genotyping Facilities at Wellcome Sanger Institute and LabCorp (Laboratory Corporation of America) using support from 23andMe. The CATSS study was supported by the Swedish Foundation for International Cooperation in Research and Higher Education (STINT), the Swedish Council for Health, Working Life and Welfare, the Söderström-Königska Foundation and the Swedish Research Council (Medicine and SIMSAM).

## Conflicts of Interest

The authors declare no conflict of interest.

## Footnotes

None.

